# From whole genome to probiotic candidates: a study of potential *Lactobacillus* strain selection for vaginitis treatment

**DOI:** 10.1101/2022.12.11.519948

**Authors:** Jinli Lyu, Mengyu Gao, Shaowei Zhao, Xinyang Liu, Xinlong Zhao, Yuanqiang Zou, Yiyi Zhong, Lan Ge, Haifeng Zhang, Liang Xiao, Xiaowei Zhang

## Abstract

Vaginitis is a syndrome characterized by not only the invasion of pathogens, but also the lack of *Lactobacillus*. *Lactobacillus* supplementation emerged as an innovative therapy for vaginitis which targeted the vaginal microbiota in recent years. However, limited live biotherapeutic products were explicitly marketed for this. The aim of our study is to characterize the potential *Lactobacillus* candidates for enhancing the treatment of vaginitis. Initially, 98 *Lactobacillus* isolates with whole genome were selected into the candidate pool. The genomic pathways involved in the biosynthesis of probiotic metabolites, adhering ability and acid/bile resistance, as well as sequences of antibiotic resistance genes, plasmids, transposons, viruses, and prophages were annotated. A scoring model was then established based on genomic analysis for ranking the performance of the strains, and the top-performing ones were applied to *in vitro* tests for sub-screening. As a result, two candidates were selected, and their probiotic abilities were verified with three reference strains. *L. crispatus* LG55-27 and *L. gasseri* TM13-16 presented outstanding ability to produce D-lactate and adhere to human vaginal epithelial cells. They also showed higher antimicrobial activity against *Gardnerella vaginalis, Escherichia coli, Candida albicans, Staphylococcus aureus* and *Pseudomonas aeruginosa* than control ones. Their acid/bile-resistant ability suggested the potential of oral supplementation. Strain LG55-27 and TM13-16 also display improved effects on pH changes, and the production of inflammation cytokines in BV model rats. This study provided a probiotic dosage recommendation for vaginitis treatment by demonstrating an effective probiotic screening pipeline based on genome sequences, *in vitro* tests and *in vivo* BV model experiment.

## Introduction

The vagina is a complicated and unique micro-ecosystem, characterized by diverse microorganisms(Buchta, 2018). Ravel et al. clustered the vaginal microbial community into five groups, four of which are dominated by *Lactobacillus* species and termed community state type (CST) I, II, III and V. The dominant species for these CSTs are *L. crispatus, L. iners, L. gasseri, and L. jensenii*, respectively(X. Zhou et al., 2010). While, CST IV is colonized by more anaerobic bacteria instead of *Lactobacillus* and exhibits higher microbial diversity. Although women with vaginal CST IV could persist this community state for a long period of time without any symptom, they are exposed to a higher risk of female reproductive diseases (Gupta, Kakkar, & Bhushan, 2019). Hence, a considerable abundance of *Lactobacillus* is regarded as the indicator of a healthy vaginal environment. The protective functions of *Lactobacillus* mainly focuse on: (1) producing the metabolites that are able to inhibit the pathogens, such as D/L-lactate, H_2_O_2,_ and bacteriocins (Tachedjian, Aldunate, Bradshaw, & Cone, 2017); (2) releasing lactic acid to keep a low vaginal pH (pH ≤ 4.5) via acidifying the vaginal environment by fermenting glucose (Wilson et al., 2010) (3) maintaining the vaginal mucous membrane barrier by adhering to the vaginal epithelial cells; (4) regulating mucosal immune response by producing short-chain fatty acids (SCFAs) to alleviate inflammatory injury (Aldunate et al., 2015).

The lack of *Lactobacillus* species in the vaginal microbiota and the presence of pathogens, such as *Gardnerella vaginalis, Escherichia coli* and *Candida albicans*, will both lead to the dysbiosis of vaginal microbiota, which results in a higher risk of vaginitis. Vaginitis is one of the most common gynecological diseases that occur frequently in women of reproductive age, including bacterial vaginosis (BV), aerobic vaginitis (AV) and vulvovaginal candidiasis (VVC), etc. They are directly related to many serious complications, such as preterm birth, postpartum endometritis, cervicitis, and pelvic inflammatory disease (Carr, Felsenstein, & Friedman, 1998). Additionally, positive treatment of vaginitis can reduce the risk for acquiring sexually transmitted diseases like human papillomavirus (HPV) and human immunodeficiency virus (HIV) infection. Antibiotic intervention, a traditional therapy, is the current clinical standard of care (SOC) for vaginitis during decades (Oduyebo OO, 2009). Although the short-term cure rate of metronidazole therapy for BV treatment was up to 80%-90%, the recurrence rate appeared as 52% after 6 months of follow-up and 68% after 12 months. Prolonged metronidazole treatment period cannot reduce the recurrence rate after drug withdrawal but increases the incidence of VVC(Bagnall & Rizzolo, 2017). The traditional antimicrobial azole derivatives can cure the most of primary acute VVC, but failed in preventing recurrent attacks(Kang, Han, Kim, Paek, & So, 2018). As a result, long-term treatment with antibiotics cannot decrease the recurrence rate. This is also true as clindamycin for AV (Donders, Ruban, & Bellen, 2015). Therefore, novel treatment approach is urgently needed for radical cure of vaginitis.

Although pathogens can be cleaned by antibiotic, the number of *Lactobacillus* is still suppressed after treatment, which may be the main reason for the high recurrence rate after SOC(Sobel et al., 2006). Hence, the supplementation of *Lactobacillus*-containing probiotics adjunct to antibiotic treatment has been developed as a novel biotherapy, and has shown beneficial effects, although heterogeneous outcomes presented in clinical trials. It is hard to achieve a probiotic *Lactobacillus* strain that could be widely effective for all vaginitis patients, because the *Lactobacillus* strains colonized in vagina are varied among different ethnic women. What’s more, few *Lactobacillus* strains were developed as live biotherapeutic products (LBPs) explicitly marketed for female reproductive health. Hence, more potential *Lactobacillus* strains should be discovered.

In this study, we aimed to select the potential female probiotic strains from a pool of *Lactobacillus* strains from a human commensal bacteria collection constructed by BGI-Research, China. Based on the whole genome of the strains, a high throughput pre-screening pipeline for probiotic was built. Subsequently, an *in vitro* test profile of preferred product characteristics, including growth rate, probiotic metabolites producing ability, adhesion to vaginal epithelial cells, gastral acid/bile tolerance, antibiotic resistance, and antimicrobial capacity, was established and carried out to further identify the best-performing isolates.

## Methods

### *Lactobacillus* isolation, culturing and sequencing

Samples from human body-site were collected in 2 ml Self Standing Screw Cap Tubes (Axygen) with storage buffer (MGI, China), and then gradient diluted in phosphate-buffered saline (PBS), spread on sterile de Man, Rogosa, Sharpe (MRS) medium, and incubated at 37°C under the anaerobic condition for 24 hours. Individual colonies were picked based on different morphology and transferred into MRS broth. After 24 hours anaerobic incubation at 37°C, 2ml culture was sent for whole genome sequencing to Illumina Hiseq 2000 platform. The whole genomes and taxonomic information were obtained as previous described(Zou et al., 2019).

All strains belonging to *Lactobacillus* species were selected to build our candidate’s pool. The scaffold sequences of the *Lactobacillus* isolates were annotated against 7 metabolism databases to comprehensively choose the suitable strains, which includes the genes involved in the metabolic pathway related to acid resistance, bile resistance, lactate producing, hydrogen peroxide producing, butyrate synthesis, propanoate synthesis, adhesive property. The vaginal dominate species were regarded as the key objects of selection. In addition, the Virulent Factor Database (VFDB) (B. Liu, Zheng, Jin, Chen, & Yang, 2019) and the Comprehensive Antibiotic Research Database CARD (Alcock et al., 2020) were also employed to investigate the safety of strain for human use. The databases were originally built from KEGG (Kanehisa & Goto, 2000), in which the enzymes were selected based on KO information. BLAST (Altschul, Gish, Miller, Myers, & Lipman, 1990)was used in the annotation process, and the e-value was set as 0.01, while identity value should be greater than 60%. In order to improve the selection quality, we used subject length to calculate the coverage of aliment length, the coverage greater than 80% and the length of aliment length larger than 100 bp (Amino acid) should be remain. A score system was established after all genomes of the *Lactobacillus* isolates were blasted against selected datasets. The highest numbers were scaled to 100 as the standard, rest numbers were standardized based on this scale. Then, the weight was added to different functions.

### Assessment of growth

1 ml 1.7 x 10^6^ CFU of preselected candidate strains was added into fresh MRS broth in triplicates under anaerobic condition at 37 □ in 96-well plates for 24 hours. The growth state of strains was determined by measuring the density of cultures at a wavelength of 595 nm (OD_595_) at 0, 2, 4, 5, 7, 9, 12 and 24 hours and the growth rate was calculated based on the logarithmic phase of the growth curve.

### Determination of D- and L-lactate production

The candidate isolates were cultivated at 37 □ in MRS broth for 48 hours in an anaerobic chamber. Then they were centrifuged at 8000 rpm for 5 min. Supernatants were collected. The concentrations of D-/L-lactate were measured by D- /L-lactic acid Assay Kits according to the manual assay procedure (Megazyme) using ultraviolet spectrophotometry. Experiments were performed in triplicates.

### Determination of hydrogen peroxide production

100 μl of each isolates incubated at 37□ for 24 hours was plated on MRS agar with 0.25 mg/mL tetramethyl benzidine (TMB) and 0.01 mg/mL horseradish peroxidase (HRP) and incubated at 37 □ for 48 hours under anaerobic conditions (Tomás, Bru, & Nader-Macías, 2003). After exposure to ambient air, colonies turn blue within 0 to 30 min depending on the concentration of hydrogen peroxide.

The candidate isolates incubated in MRS broth for 48 hours were centrifuged at 8000 rpm for 5 min. The supernatants were collected and the concentration of H_2_O_2_ was measured by the Hydrogen Peroxide Assay Kit (Nanjing Jiancheng Bioengineering Institute) according to the manual assay procedure. Experiments were performed in triplicates.

### Evaluation of auto-aggregation

The auto-aggregation ability of *Lactobacillus* strains was evaluated by the spectrophotometer at 595 nm (OD_595_). 10 mL of 10^8^ CFU/mL *Lactobacillus* strains cultured for 24 h was added in a 15 mL test tube. The upper suspensions were tested separately at 0 min, 30 min, and 60 min. Auto-aggregation was assessed by following equation: auto-aggregation rate (%) = [1 - (OD_60min_/OD_0min_)] × 100 (Angmo, Kumari, Savitri, & Bhalla, 2016).

### Adhesion ability to vaginal epithelial cells VK2/E6E7

The vaginal epithelial cells VK2/E6E7, with a growth density of up to 80% in complete DMEM medium (Gibco), were digested with trypsin and resuspension in fresh medium at a concentration of 2 ×10^5^ cells/ml.1ml suspension was added to each well in 6-well plates and incubated at 37 □ in a 5% atmospheric CO_2_ environment until the cells were grown to 80% confluence. Cells were then washed twice with PBS, and 1 ml fresh liquid medium was added into each well.

*Lactobacillus* strains were cultured in liquid MRS for 24 hours at 37 □. Then 1 mL suspension was added into 6-well plates with VK2/E6E7 and co-incubated at 37 □ for 90 min in 5% CO_2_. Cells were washed five times with 1 ml PBS to remove unbound bacteria and 1ml trypsin was added into well to cause cell lysis. DMEM medium with Fetal bovine serum was then added. 1 mL of 0.05% Triton X100 was added for 5 min. Ten-fold gradient dilution was applied, and the culture was plated on MRS agar. Colonies were counted after 24 hours incubation at 37 □ under anaerobic conditions. Cell counts were performed by automatic cell counter.

### Antimicrobial activity testing

*Lactobacillus* strains were incubated in MRS broth for 24 hours under an anaerobic condition. The suspension was filtered using a 0.2 μm filter membrane when *Lactobacillus* cells number reached 1 **×** 10^8^ CFU/ml. 100 μl of *G. vaginalis, E. coli, C. albicans, S. aureus, P. aeruginosa and E. cloacae* suspension incubated in MRS broth for 24 h was added into 1.5 ml *Lactobacillus* supernatant. After 24 hours of co-incubation, absorbance at OD_595_ was then measured. The same procedure was carried out for *E. coli* and *C. albicans*.

### Antibiotic susceptibility testing

Kirby-Bauer test (including sixteen antibiotics) was applied to evaluate the antibiotic susceptibility. 100 mL of *Lactobacillus* culture was incubated for 24 h at 37 □ under anaerobic condition. Then it was spread on the MRS agar. Antibiotic disks were put on to MRS medium. After 24 hours, the radius of inhibition zone was measured.

### Acidic and bile salt tolerance measurement

To investigate the influence of low pH on *Lactobacillus* growth, 100 ul of *Lactobacillus* suspension incubated at 37□ for 24h under anaerobic environment was added into MRS broth with pH = 2, 3, 4, 4.5 and 7. Then the culture was plated on MRS agar after 24 hours of incubation. Colony counting was then performed. The same process was repeated to access the influence of high concentration of bile salt on the growth of *Lactobacillus* strain using MRS broth with 0.05%, 0.1%, 0.2% and 0.3% of bile salt.

### Administration of *lactobacillus* strains on BV model rats

#### Modeling

Seven-week-old female SD rats, weighed 222-264 g, were obtained from Hunan SJA Laboratory Animal Co. (Hunan, China). A total of 220 female rats were examined during the experiment period. The rats were housed in dry, stainless-steel cages that contained 2-5 rats each, and they were cultivated in a standard animal maintenance facility under a 12h/12h light/dark cycle at 20.0-24.4□ and 40-70% humidity. The experiment was performed under pathogen free conditions in order to avoid unwanted infection.

0.5 mg estradiol benzoate purchased from Shanghai full woo Biotechnology (Zhumadian) Co., Ltd. (Shanghai, China) was injected into rats three days before *G. vaginalis* infection in order to maintain the rat pseudoestrus. Then, 150 μl 1×10^8^ CFU/ml *G. vaginalis* were injected vaginally once a day for three days. The clinical characteristics, including hyperaemia, oedema, infiltration and bleeding were observed after infection.

### Grouping and treatment

The successful modeled rats were randomly divided into 16 groups of 10 rats per group (Figure 1). The Model group was set as control with no treatment; the Metronidazole group was treated with metronidazole intragastrically (10 mL/kg body weight each day) for 2 days after modeling and without treatment in the following days. As described in Figure 5, the remaining groups were administrated with bacteria both intragastrically (I) and vaginally (V), each of which contained 7 groups: LG55-27 (LG, administrated with 1×10^8^ CFU/day for 6 days), TM13-16 (TM, administrated with 1×10^8^ CFU/day for 6 days), low dose of LG55-27+TM13-16 (LC, administrated with 1×10^4^ CFU/day for 6 days), medium dose of LG55-27+TM13-16 (MC, administrated with 1×10^6^ CFU/day for 6 days), high dose of LG55-27+TM13-16 (HC, administrated with 1×10^8^ CFU/day for 6 days), high dose of LG55-27+TM13-16 and metronidazole (HCM, administrated with metronidazole and 1×10^8^ CFU/day probiotics on day 0-2, followed by probiotics alone with 1×10^8^ CFU/day on day 3-6), and inactivated LG55-27+TM13-16 (SC, administrated for 6 days).

**Figure 1.**
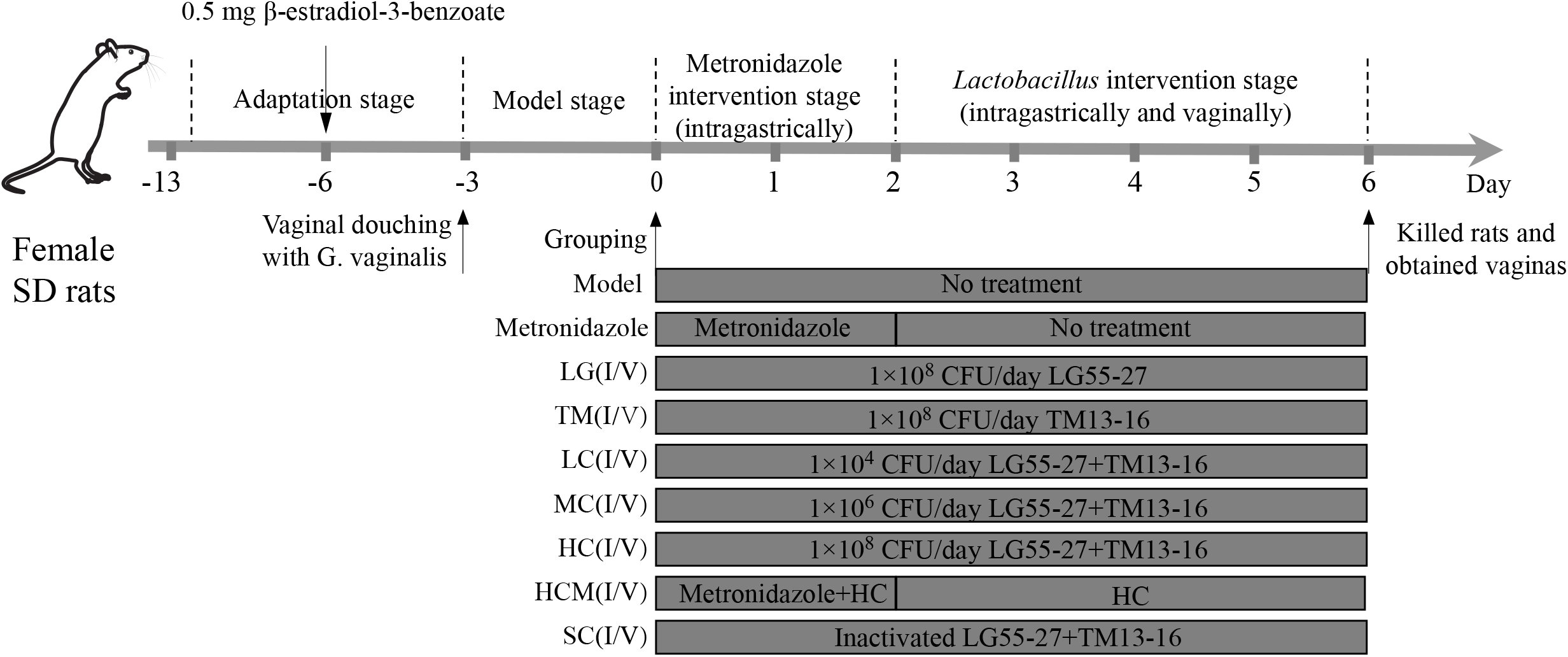
The process of intervention of rats and grouping after BV model was established.

Metronidazole Tablets (Shanghai Sine Wanxiang Pharmaceuticals Co., Ltd., China) were dissolved in pure water, making a concentration of 0.0108 g/mL. The pellet of strains was resuspended by PBS and prepared into different concentration respectively. The mixed strains were prepared with total concentration of 1×10^8^ CFU (radio of 1:1). The inactivated strains were autoclaved at 121 □ for 15 minutes before administration.

### Detection of inflammation cytokines, pH value and colonization of *Lactobacillus*

On day 0,3,5, the pH value of the rats’ vagina was measured. The vaginal discharge and the fecal sample were collected on day 0 and 5. Genomic DNA was extracted (TIANamp Bacteria DNA Kit, TIANGEN Biotech, China) and examined the amount of *L. crispatus* and *L. gasseri* using primers R-GV3 (CCGTCACAGGCTGAACAGT) and LcrisF (AGCGAGCGGAACTAACAGATTTAC), LcrisR (AGCTGATCATGCGATCTGCTT) and LactoF (TGGAAACAGRTGCTAATACCG), respectively(De Backer et al., 2007). Rats was killed on day 6 and the vaginal tissues were collected and homogenized. The cytokine levels, including IL-1β and TNF-α, were assessed using the Elisa Biotech cytokine Kits (Shanghai Elisa Biotech Co., Ltd., China) according to the instruction.

## Results

### Pre-screening based on whole genomes of *Lactobacillus* strains

In total, a *Lactobacillus* pool containing 98 strains isolated from various human body sites of different participants were prepared(Zou et al., 2019). Six functional gene datasets were prepared based on KEGG in-house, which includes the genes involved in the metabolic pathway related to acid resistance, bile resistance, lactate production, hydrogen peroxide production, butyrate synthesis, propanoate synthesis and adhesive property. In BLAST results, the sum of the numbers of enzymes annotated by each strain was calculated as one of the factors in our score system. The function in the right part of figure 1 gains more weight than left, due to they considered to contribute more in environment in future (Supplementary Table 1). The score of safety was individually calculated because we believe it provides negative contribution, thus, they were given negative weight in our score calculating process. 23 *Lactobacillus* strains were preliminary according to the final score shown in the last column of Figure 2.

**Figure 2.**
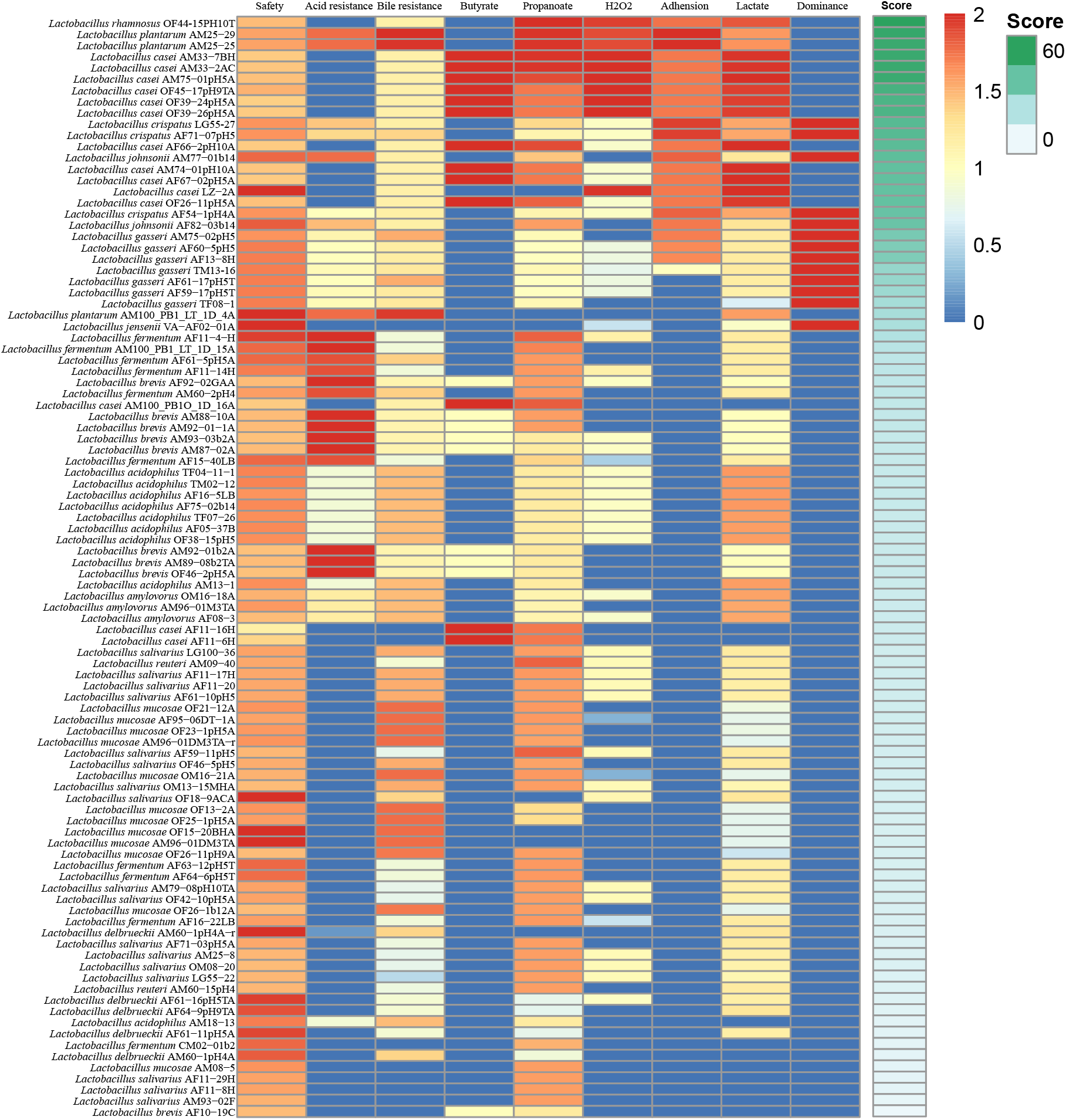
*Lactobacillus* strains were selected based on the whole genome annotation results.

### Further screening based on *in vitro* tests of pre-selected *Lactobacillus* strains

D-/ L-lactate and hydrogen peroxide production, growth rate was tested (Figure 3). The growth rate of LG55-27 was the highest among all strains, which was significantly higher than AF71-01pH5 with the lowest growth rate (P < 0.0001). AM77-01b14 and TM13-16 were the highest among *L. jensenii* and *L. gasseri* separately. OF45-17pH9TA was the lowest among the *L. casei* species (Figure 3a).

**Figure 3.**
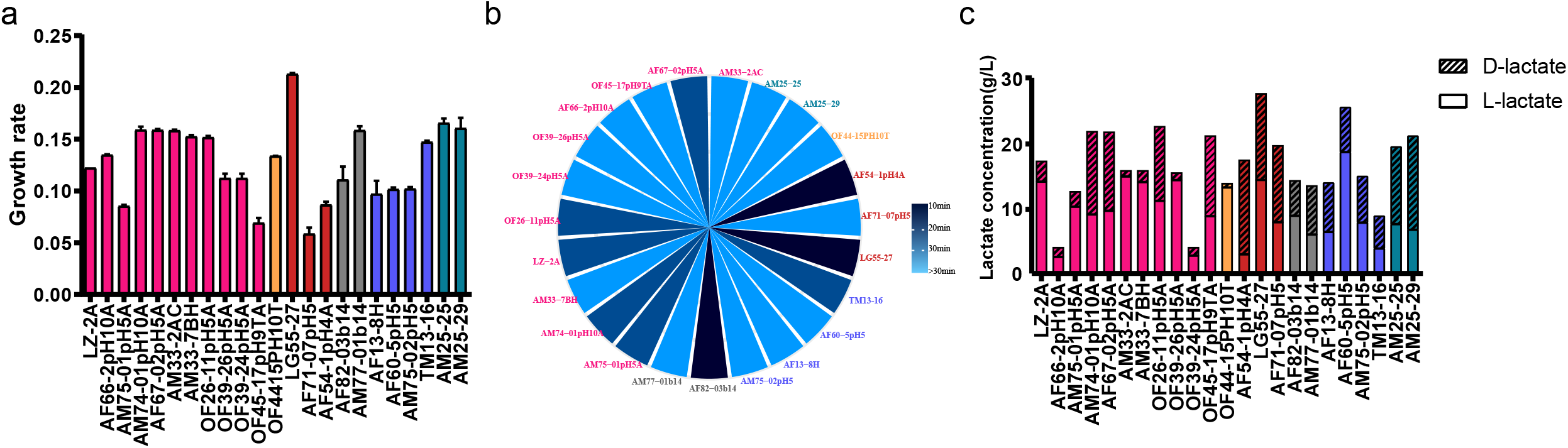
In- vitro tests were applied to screen qualified *Lactobacillus*.

The production of hydrogen peroxide was tested by semi-qualitative TMB-HRP assay. The colony of LG55-27, AF54-1pH4A and AF82-03b14 turned blue in 10 minutes, suggesting higher production of H_2_O_2_, while the colony of the other 14 isolates didn’t turn blue in 30 minutes, showing the ability of H_2_O_2_ production.

D-/L-lactate is considered as the most crucial characteristic of vaginal Lactobacillus, which provides an acidic environment in vagina and has anti-microorganism activity(Borges, Silva, & Teixeira, 2014). Therefore, the ability to produce D-/L-lactate was important when screening the candidate strains. The results were shown in Figure 3c. The ability to produce D-/L-lactate had great disparity among different strains of the same species. The total lactate (D- and L-) produced by LG55-27 was the highest among all of the species. Furthermore, LG55-27 produced the same level of both L-lactate and D-lactate, which is optimal for *Lactobacillus*. Some strains produced high level of L-lactate but almost no D-lactate. In contrast, some strains produce high level of D-lactate and low level of L-lactate (such as AF54).

*L. crispatus, L. iners, L. gasseri*, and *L. jensenii*, were characterized as the highest abundance of lactobacillus in vagina(X. Zhou et al., 2010). *L. rhamnosus* was chosen to treat BV in recent research studies (Chee, Chew, & Than, 2020). Taking these 5 lactobacilli and the result of growth rate and production of H_2_O_2_ and D-/L-lactate into account, LG55-27 (*L. crispatus*) and TM13-16 (*L. gasseri*) were screened as candidates for the subsequent functional evaluation *in vitro*. Three strains, *L. reuteri* RC-14, *L. rhamnosus* GR-1 and *L. delbrueckii* DM8909 isolated from 2 commercial vaginal probiotics found on the Chinese market were chosen as control.

### Functional validation of the candidate probiotic strains

#### Antimicrobial properties

Supernatants from the culture of five *Lactobacillus* strains were used to evaluate the ability to inhibit the growth of five typical pathogens (*G. vaginalis, E. coli, C. albicans, S. aureus* and *P. aeruginosa*). The growth of *C. albicans* was evaluated under aerobic conditions. And the growth of other pathogens was evaluated under anaerobic conditions. All five *lactobacillus* strains above significantly inhibited *C. albicans*, E. *coli, S. aureus and P. aeruginosa* (Figure 4a). All strains significantly inhibited *G. vaginalis* with the exception of RC-14 (P=0.0603). LG55-27 and TM13-16 tended to have a better antimicrobial ability to inhibit *C. albicans* than reference strains. All five *Lactobacillus* showed similar antimicrobial performance against *E. coli, S. aureus* and *P. aeruginosa*

**Figure 4.**
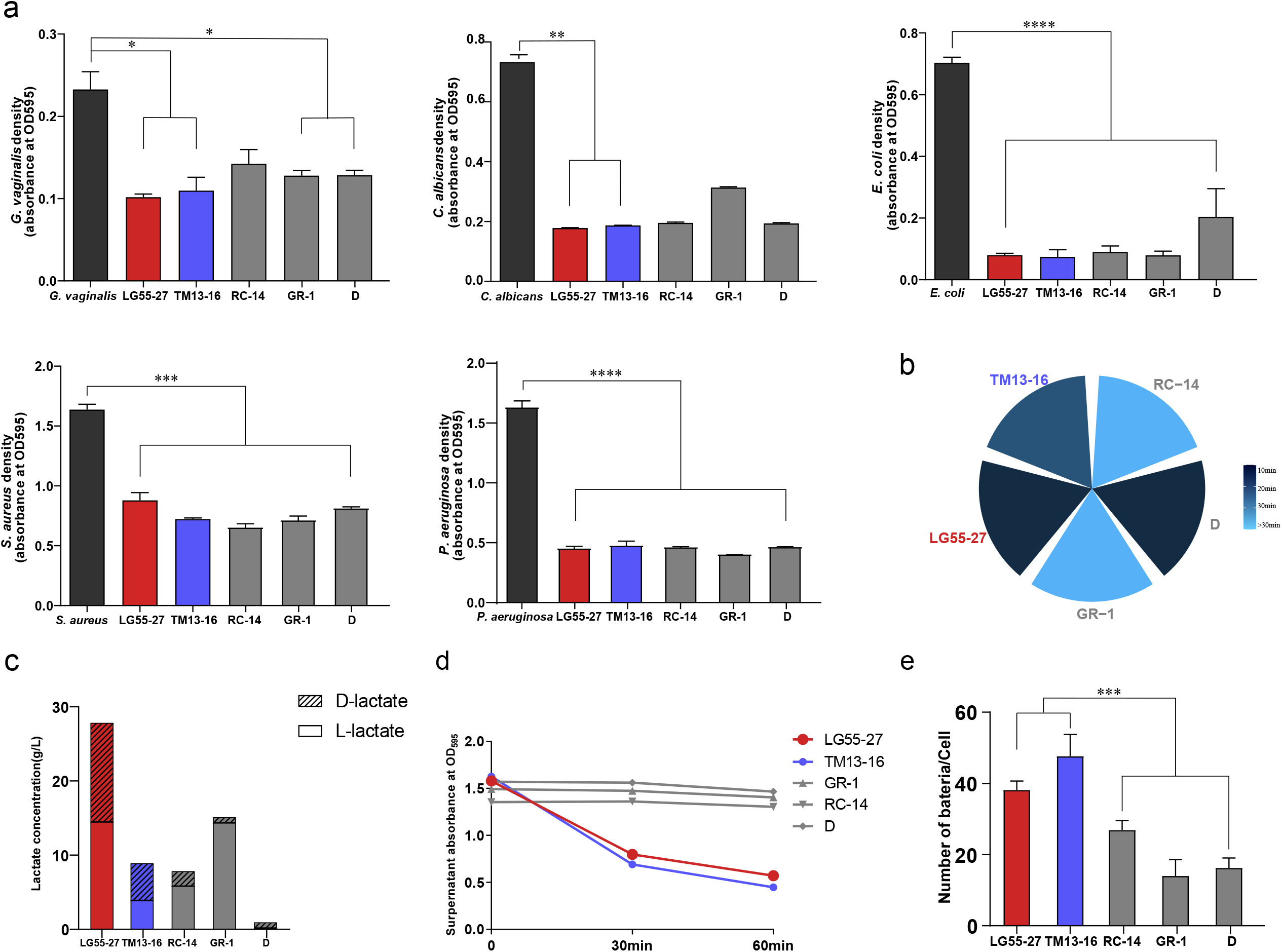
Functional verification of candidate strains. (a) Antimicrobial characteristics of *Lactobacillus* strains. (bc) Producing ability of H_2_O_2_ and L-/D-lactate for candidate strains and reference strains. (d) The ability of self-aggregation. The supernatant was measured by microplate reader at 0,30 and 60 min. (e) Adhesion ability of candidate strains and reference strains to vaginal epithelium VK2/E6E7 cells. Statical analysis were performed with Kruskal-Wallis test. * stands for *p*-value < 0.05; ** stands for *p*-value < 0.01; *** stands for *p*-value < 0.001; **** stands for *p*-value < 0.0001.

### Measurement of hydrogen peroxide

The production of hydrogen peroxide was tested by semi-qualitative TMB-HRP assay. LG55-27 and D isolates produced the most H_2_O_2_, which isolates turned blue in 10 minutes. The other species evaluated produced lower amounts of H_2_O_2_ (Figure 4b). TM13-16 turned blue in 20 minutes, producing moderate H_2_O_2_. GR-1 and RC-14 didn’t turn blue in 30 minutes, exhibiting low ability to produce H_2_O_2_.

### Measurement of D-/L-lactate production

The production amounts of D-/L-lactate produced by the two candidates Lactobacillus and three references were evaluated. LG55-27 produced high levels of both D-/ L-lactate (Figure 4c), especially D-lactate compared to other isolates (P<0.0001). GR-1 produce a higher concentration of L-lactate comparing to D-lactate production. The concentration of L-lactate produced by 2 candidate *Lactobacilli* was significantly higher than the reference strains except RC-14 (LG55-27 vs RC-14: p<0.0001, LG55-27 vs D: p=0.0006, TM13-16 vs RC-14: p<0.0001, TM13-16 vs D: p=0.0023,). The pH of the supernatant of isolates was also measured (Supplement Figure1). The ability of strains to lower the culture pH was negatively correlated to the concentration of L-lactate and D-lactate (Spearman rho = -0.457, p = 0.158 for L-lactate; Spearman rho =0.309, p = 0.355 for D-lactate), indicating that the lactate produced by *Lactobacilli* may account for vaginal acidity.

### Self-aggregation

Self-aggregation, a ability of probiotic bacteria to form cellular aggregates, is considered a desirable property, as lactobacillus can potentially inhibit adherence of pathogenic bacteria to epithelial by forming a barrier(Goh & Klaenhammer, 2010). TM13-16 and LG55-27 showed a significantly higher aggregation level than other strains after 1-hour incubation. LG55-27-16 and TM13-16 formed clumps that sedimented in a short time, leaving the supernatant clear after 30 minutes. In the first 30 minutes, the self-aggregation level of TM13-16 and LG55-27 was higher than 30 to 60 minutes. The decrease in absorbance for all strains was shown in Figure 4d. LG55-27 and TM13-16 showed a significant decrease in OD_595_ (p <0.0001), and there were no significant changes among the other strains.

### Adhesion to vaginal epithelial cells

It is an essential step when *lactobacillus* adherent to vaginal epithelial cells as it aids in its colonization and inhibition against the pathogen. Thus, the adhesion ability of *lactobacillus* to vaginal epithelial cells VK2/E6E7 was evaluated *in vitro*. It was observed that TM13-16 displayed the highest adhesion capacity to VK2/E6E7 among all strains, and LG55-27 and TM13-16 were significantly higher than other strains (p <0.01). The high level of self-aggregation of TM13-16 may hold responsible for its high adhesion ability (Spearman rho = 0.759, p < 0.01). There were no statistically significant differences in adhesion abilities among the rest of the strains (Figure 4e).

### Resistance to gastral acid and bile salt

The *Lactobacillus* growth was negatively influenced by the decrease in medium pH, leading to a reduction of live *Lactobacillus*. All strains showed a steady loss in viability when incubated at 37□ for 24 hours (Figure 5a). Initially, the viability of the two strains was 10^8^ CFU/ml at pH 7.0. LG55-27 decreased to 10^6^CFU/ml after 24-hours incubation, while TM13-16 further decreased to 10^4^ CFU/ml, indicating that it is the most acid-sensitive strain.

**Figure 5.**
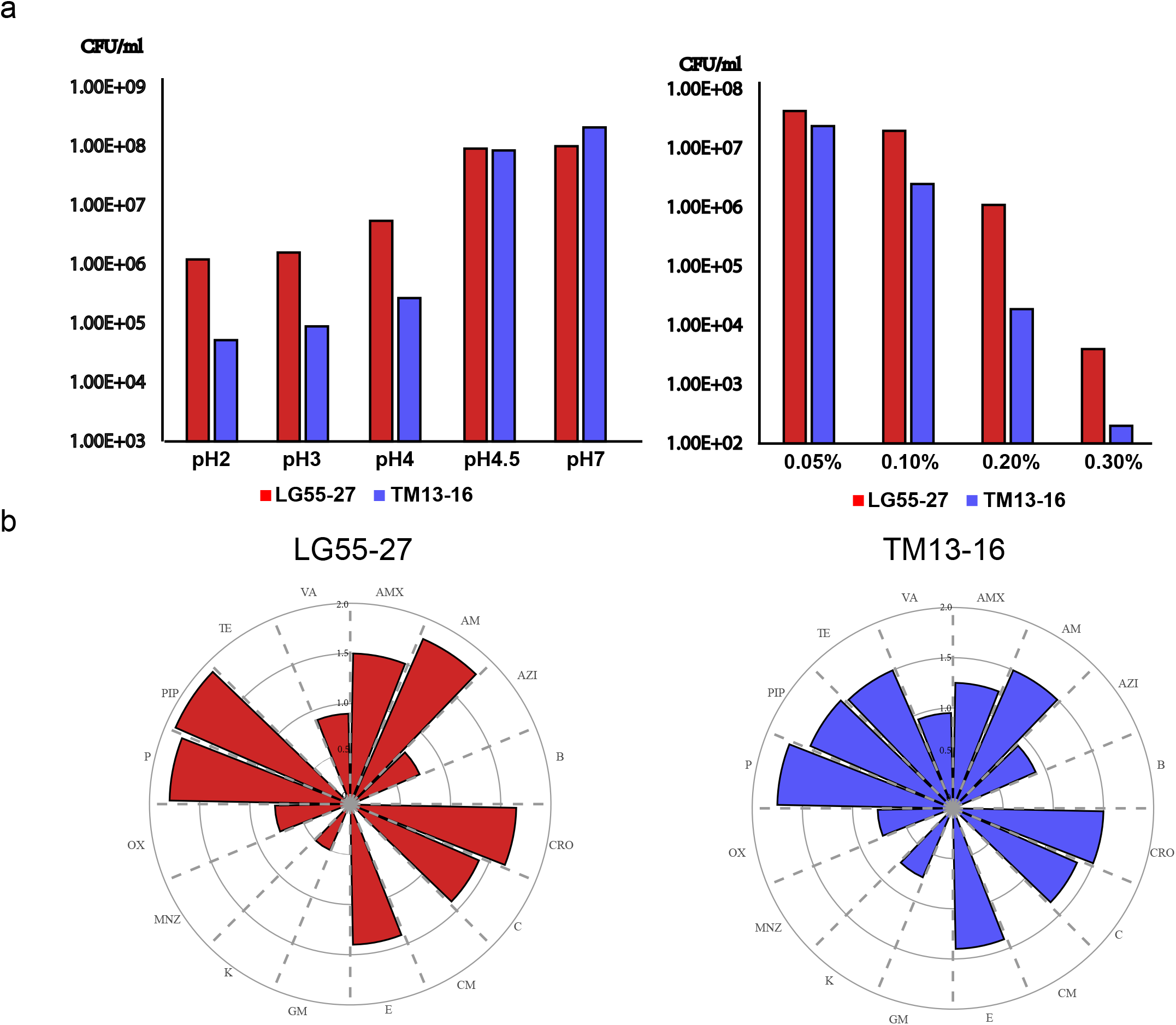
Growth ability of strains in extreme conditions. (a)Growth ability of candidate strains under different pHs and different concentration of bile salt. (b) Antibiotic susceptibility of candidate strains.

The growth of *Lactobacillus* was positively correlated with the concentration of bile salt in the medium. All strains showed a loss of viability after 24-hours of incubation at 37□. Similar to the result of the low pH resistance test, the viability of TM13-16 was reduced the most compared to other strains. LG55-27 reduced to 10^3^CFU/ml and from 10^7^ CFU/ml. All strains exhibited a strong survival ability in MRS under the anaerobic condition.

### Antibiotic resistance

We characterized phenotypic antibiotic resistance profiles of the 2 lactobacillus strains to 16 kinds of antibiotics. All of them revealed phenotypic susceptibility to many antibiotics, including Penicillin, Ampicillin, Piperacillin, and so on (Figure 5b). Two of them showed similar susceptibility to Penicill, Piperac, Erythrocin, and Chloramphenicol. TM was the most susceptible to Tetracycline, while LG55-27 was the least. All of them were resistant to Bacitracin and Metronidazole. LG55-27 was resistant to tetracycline. TM13-16 was Clindamycin and Gentamicin resistant.

### Safety analysis of select *Lactobacillus* strains

The scaffold of two selected strains, TM13-16 and LG55-27 were used for safety analysis. We use the scaffolds to perform with the CARD, Resfinder, the VFDB database, prophage, and plasmids, as well as the Mobile genetic elements databases.

The antibiotic resistance of two strains was predicted using CARD (Alcock et al., 2020) and Resfinder (Zankari et al., 2012). The annotation of CARD database shows the tetracycline antibiotic and macrolide antibiotic genes in the strain LG55-27, carbapenem, lincosamide antibiotic, only tetracycline antibiotic in TM13-16. The results of Resfinder illustrate the Macrolide resistance and Tetracycline resistance of LG55-27, and tetracycline antibiotic of TM13-16.

The potential prophage sequences were identified using PHAST (Y. Zhou, Liang, Lynch, Dennis, & Wishart, 2011) database. None of the two strains contain the prophage gene under the threshold identity>90%, e-value >0.01 and minimum aligned length>5k bps. The mobile genetic elements were also analyzed using blast search against the corresponding database. Islander (Hudson, Lau, & Williams, 2015) was utilized to analyze Genomic island, and ICEberg (M. Liu et al., 2018) database was performed to identify the Integrase and conjugative elements. The insertion sequence was annotated against Isfinder (Siguier, Perochon, Lestrade, Mahillon, & Chandler, 2006) database. We did not find any genomic island or integrase and conjugative elements in the scaffolds of our two strains. For the result of insertion sequences, the annotation demonstrates 2 hits in TM13-16. Those hits exist in different scaffolds and may not be capable to trigger the transposition activity. In the strain LG55-27, 22 hits were found similar to insertion sequences. Although some of the hits are located on the same scaffold, possibly leading to transposition activity, we believe there are no harmful genes in LG55-27 to be transposed. In the results of Virulence factors, the cut off was e-value >0.01, coverage >70%, identity>30%. Under this threshold, 151 potential virulence? genes were identified in TM13-16, and 175 potential virulence?gens were identified in LG55-27. The annotated function mainly includes secretion and transport system, structure-forming systems like capsule and cell wall, regulation molecules, cell surface proteins for better adherence and motility, and stress protein. most of these genes are defensive or non-classical virulence factors, helping improve the cell survivability in a complex environment. The two strains contain genes encoding adherence and lipopolysaccharides, which helps cells’ colonization. The toxic gene lpxM (Xu et al., 2013) did not found in their scaffolds. Notably, we found cytolysin/hemolysin related genes in all two strains. The product of this gene is a pore-forming structure, it may cause infections in humans and animals (Koronakis & Hughes, 2002). However, it is a kind of bacteriocin, harmful to a broad range of gram-positive bacteria, maintaining the balance in the intestinal environment.

### Evaluation of the probiotic administration effect on BV rats

#### *Lactobacillus* was colonized in the vagina and gut

The colonization capacity of the *Lactobacillus* was evaluated by qPCR. For the intragastrical administration groups, the abundance of the intervention strains was measured in fecal samples (Figure 6a). The results showed that the abundance of strain TM13-16 in TM and HC groups was significantly increased after 5 days of intervention, but no statistical significance showed in strain LG55-27 after the intervention. It indicated that the colonization ability of TM-13 in the intestine was higher than LG55-27. In addition, vaginal discharge was used to evaluate the colonization of LG55-27 and TM13-16 in vaginal lavage groups (Figure 6b). After intervention, the abundance of LG55-27 was significantly increased in LC, MC and HC group, and TM13-16 was significantly increased after intervention in all groups. This demonstrated a better colonization capacity of these two strains in vagina than in intestine.

**Figure 6.**
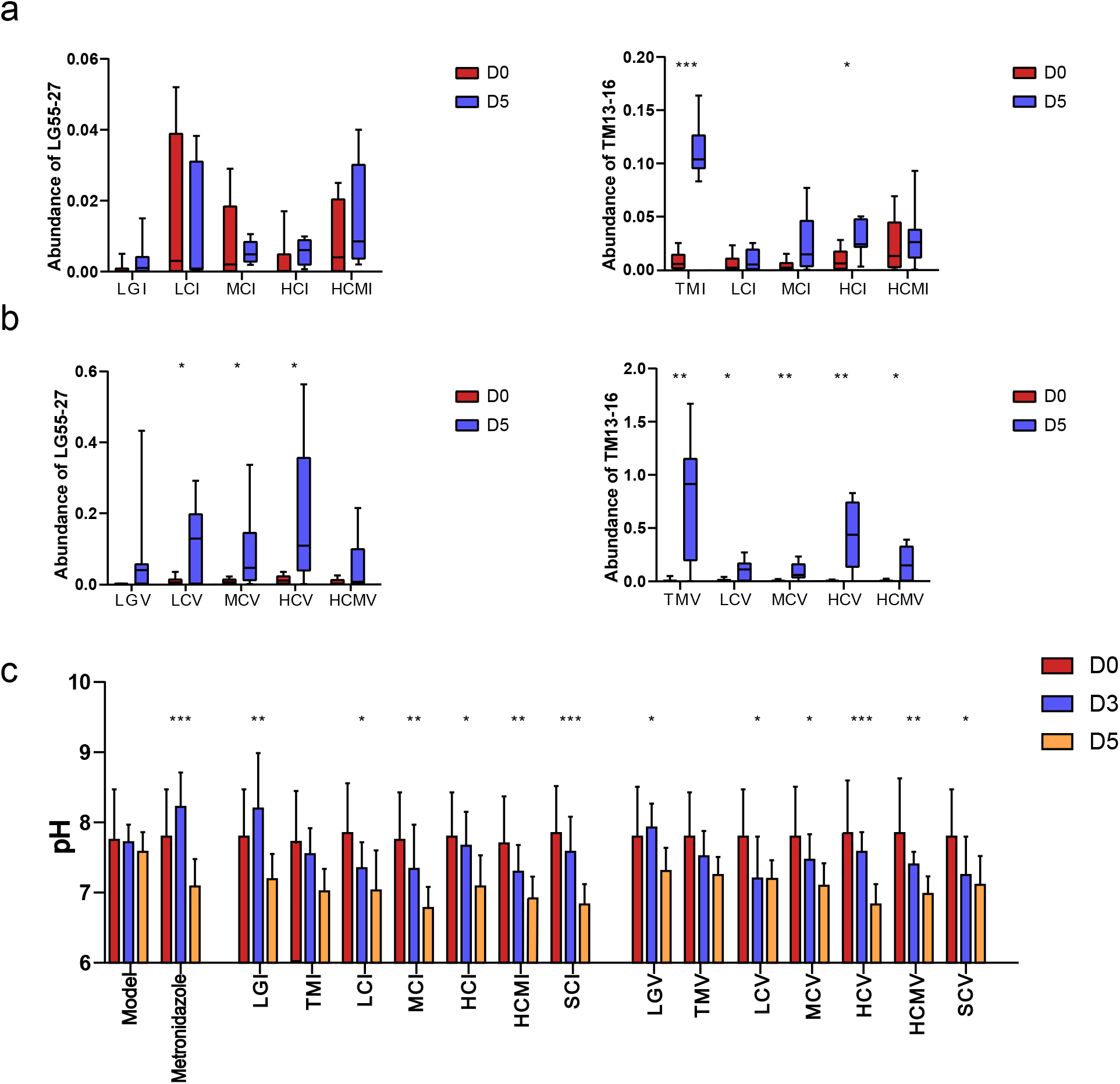
The outcomes after oral and vaginal administration. The abundance of LG55-27 and TM13-16 in fecal (a) and vaginal (b) microbiota. D0 and D5 represent the fecal and vaginal samples collected at 0-day and 5-day respectively. (c) pH value in vagina was tested at 0-day, 3day and 5day. Wilcoxon test was used to do the statistical analysis of each two independent groups. * stands for *p*-value < 0.05; ** stands for *p*-value < 0.01; *** stands for *p*-value < 0.001

**Figure 7.**
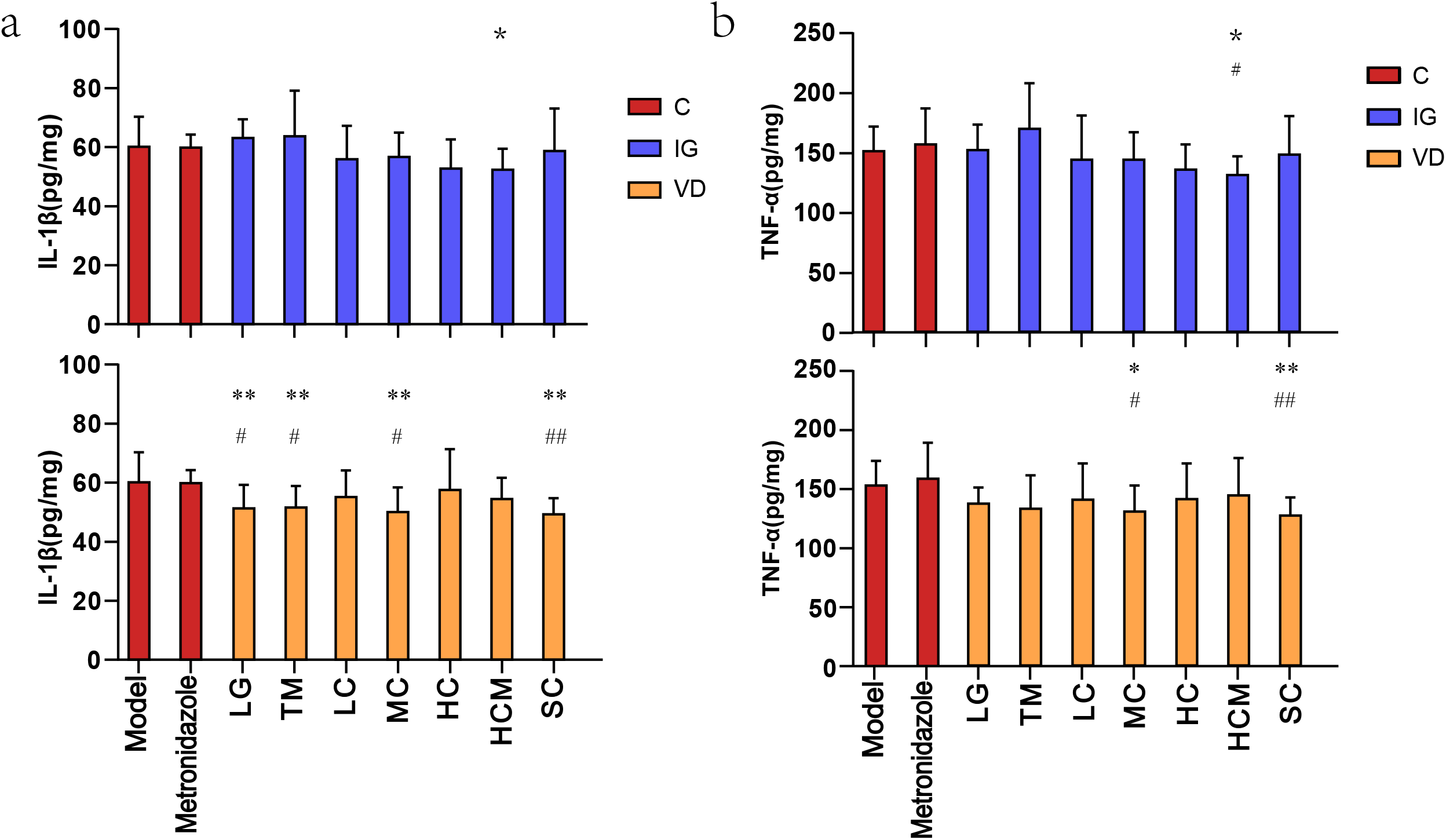
Inflammatory cytokine production of rats’ vaginal tissue in response to *Lactobacillus*. Cytokines IL-1β (a) and TNF-α (b) in vagina were measured after 6-day intervention. C represents control group; IG represents Intragastric administration group; VD represents vaginal douche group. * presents comparison to metronidazole group. * stands for *p*-value < 0.05; ** stands for *p*-value < 0.01. ^#^ represents comparison to model group; ^#^ stands for *p*-value < 0.05; ^##^ stands for *p*-value < 0.01..

#### pH changes in vagina after *Lactobacillus* intervention

The vaginal pH as a key diagnosis criteria for BV in the clinic was also monitored in BV rats after administration of *Lactobacillus* on day 0, 3 and 5 (Figure 6c). All the groups showed decreasing trend except metronidazole and LG groups on day 3, indicating a delayed effect of these two treatment methods. Furthermore, all intervention groups but TMI and TMV significantly reduced after 5 days of administration. Though the probiotics and metronidazole groups displayed considerable effect on vaginal pH value, there were no significantly different effects between groups.

#### Inflammatory cytokine responses to *Lactobacillus* in rats

Vaginal homogenate levels of interleukins (IL-1β) and tumor necrosis factor-alpha (TNF-α) were applied to evaluate the effect of *lactobacillus* after administration. For the single-strain treatment, only the vaginal lavage groups (LGV and TMV) significantly lower IL-1β, which means single strain should be more efficient by vaginal treatment than oral treatment. In addition, MCV group could significantly lower the level of IL-1β and TNF-α, indicating that medium dose of mix strain should be selected for vaginal treatment. The level of IL-1β and TNF-α were significantly decreased in HCMI group, but not in HCMV group, suggesting that probiotics adjuvant therapy of metronidazole should be treated orally. Interestingly, inactivated strains (SCV) also significantly lower the level of IL-1β and TNF-α, which demonstrated that the metabolisms of *lactobacillus* reduced the rats inflammation responses of BV. What’s more, there were no significant difference between model and metronidazole groups, implying that although metronidazole could inhibit the pathogens, it might have a delayed respond to inflammation in BV rats.

## Discussion

The high incidence and recurrence of vaginitis among women with reproductive age bring big troubles that plagued their daily life, and further lead to many severe gynecological diseases. The biotherapy with live probiotics in a combination with antibiotics has been proposed as a potential treatment method that can lower the incidence and recurrence of vaginitis. However, a lack of efficient probiotic strains for vaginal health has been developed as commercial products that functioned internationally, especially in China. According to a previous study, vaginal microbiota exhibits ethnic differences and will result in the deviation of the prevention outcomes(Antonio, Hawes, & Hillier, 1999). *L. rhamnosus* GR-1 and *L. reuteri* RC-14 are the well-developed vaginal health probiotics that have been identified in 1980s’ by Reid *et. al* and studied for decades(Reid, Cook, & Bruce, 1987). To date, plenty of clinical trials were carried out on the treatment of vaginitis and urethritis by administering the strains orally or vaginally(Ho et al., 2016; Martinez et al., 2009). Whereas, the outcomes were not consistent among participants from different countries, which appeared to be efficient in European(Cianci et al., 2008) and African (Anukam et al., 2006) but invalid in British(Jordan E Bisanz 2014), Canadian(Husain et al., 2019) and Chinese (data unpublished). What’s more, *L. delbrueckii* DM8909 are served as the active ingredient of the unique IND-approved live biotherapeutic drug around the world. These three strains are recognized as excellent probiotics for female reproductive health. Hence, we isolated them from marketed commodities as the control strains of this study. We were aimed at characterizing the *Lactobacillus* strains with better performance and wider applicability.

Traditionally, choosing a suitable strain was based on the publication, the strains with the reported function will be selected as the candidate. However, the functions may be slightly different from the publication to the actual database. Additionally, some other functions that may provide contribution could be missed from the selection process. In comparison, we build a model which includes various functions, not only the safety and prebiotics, but also the survivability. This method can be performed to the high throughput target strain selection, and rapidly pick the best results from a large quantity of data. More functions can be included in the model depending on demands. The potential of strains’ function which was not reported before are predicted at any rate.

Previous research studies in the field usually evaluated the function of bacteria with 10 or more isolates *in vitro*, which increased the complexity of the research. In this study, a genome database was established based on lactobacillus functions (producing lactic acid and hydrogen peroxide). And high-throughput screening of bacteria was performed based on the existing *Lactobacillus* isolates whole-genome database, which made the screening to be more effective and accurate. It also provides an effective method for the screening of other functional isolates. According to the current study, research associated with vagina lack of effective experimental animal models. In this research, BV model was preliminarily constructed in rats, and some clinical indicators were used to evaluate the effect of candidate strains on bacterial vaginosis. What’s more, different doses of strain were also selected to evaluate the effects, providing an experimental basis for clinical intervention.

Two candidate *Lactobacillus* strains were selected and emphatically studied in this research. Along with three control isolates from commercial products, the pH value of all these five strains cultures decreased after 24 hours of incubation (Supplement Table 1), and their supernatants all inhibited the growth of *E. coli, G. vaginalis, C. albicans, Staphylococcus aureus* and *Pseudomonas aeruginosa* significantly. These results indicate that the production of metabolisms is essential in maintaining the acid environment and forms protective barriers in the vagina. The previous study demonstrated a negative correlation between the lactic acid concentration and the pH, indicating that lactate is primarily responsible for the acidification of the vagina (Landay, O’Hanlon, Moench, & Cone, 2013). And Lactic acid produced by *Lactobacillus* is the major antimicrobial compound, hindering the growth of the *G. vaginalis* (Atassi, Brassart, Grob, Graf, & Servin, 2006), *E. coli* (Valore, Park, Igreti, & Ganz, 2002) and *C. albicans* (De Gregorio, Silva, Marchesi, & Nader-Macias, 2019) as well as other pathogens, like *Chlamydia trachomatis* (Nardini et al., 2016). D-lactate and L-lactate are the isomers of lactic acid, which is still controversial on its effects. L-lactate is normally the primary isomer of lactate in the blood of humans by metabolizing pyruvate via L-lactate hydrogenase (Petersen, 2005). D-lactate was mainly produced by bacterial fermentation in the intestinal tract and the D-lactate concentration in adults’ serum ranges from 11-70 nmol/L (Takahashi, Terashima, Kohno, & Ohkohchi, 2013). D-lactic acidosis occurs when D-lactate accumulates massively. However, L-lactate play an essential role in woman’s health. The production of D-lactate by lactobacilli enhances protection against microbial invasion of the upper genital tract(Donders, 2015). And D-lactate might have superior protective properties compared to L-lactate, although the lactate isomers have similar HIV-1 viricidal and bactericidal activity against BV-associated bacteria(Aldunate et al., 2013). An increasing ratio of D- to L-lactate correlated significantly to a decreasing concentration of EMMPRIN (Extracellular metalloproteinase inducer), indicating that MMP-8(matrix metalloproteinase) was also influenced as MMP-8 was induced by EMMPRIN (Witkin et al., 2013).The levels of EMMPRIN and MMP-8 were elevated in women with VVC (Beghini, Linhares, Giraldo, Ledger, & Witkin, 2015). The D/L-lactate ratio of 1 might actually be quite favorable(Verstraelen, Vervaet, & Remon, 2016), which is close to what is observed with *L. crispatus*. In this study, three out of five strains (LG55-27, TM13-16, GR-1) produced high levels of L-lactate with no significant differences. However, the production of D-lactate was highly varied *L. crispatus* LG55-27 produced the highest level of D-lactate, and it was significantly higher than other strains. The D/L-lactate ratio of LG55-27 and TM13-16 were 1.00±0.11 and 1.16±0.19 respectively, which were suitable for woman health. The D/L-lactate ratio of other strains was far from 1, indicating that the LG55-27 and TM13-16 were more favorable to be the candidate probiotics. Since the complex formula is commonly applied in the marketed probiotic production to make up for the deficiency of single strain, the concentration of D-lactate and L-lactate produced by a mixed culture of LG55-27 and TM13-16 was also evaluated (Supplement Figure 1). The total lactate (D-lactate and L-lactate) production of these combinative strains was significantly higher than any of the single strains and also higher than the combination of two reference strains GR-1 and RC-14. Therefore, the compound of the two candidate strains holds immense potential to be promoted as an LBP for vaginal health.

Studies have shown that both topical and oral probiotics can improve the microenvironment of the vagina. Under the condition of vaginal probiotic treatment, strains may effect by adhering to the vaginal epithelial cells, and further produce lactic acid, hydrogen peroxide and antibacterial metabolites. However, it is demonstrated that the vaginal microbial strains are individual-specific, and the native *lactobacilli* colonized persistently, while the foreign ones colonized transiently (A C Vallor, 2001). Even so, the exogenic lactobacilli improve survival rate and promote the growth of the native vaginal microbiota, thus supporting the natural barrier against pathogenic microorganisms(Borges et al., 2014). Therefore, the adhesion of the *Lactobacillus* strain to the human vaginal epithelia cells is a crucial factor for its effect. TM13-16 and LG55-27 significantly adhered to the vaginal epithelia VK2 compared to other strains, competing with pathogens for adhesion sites, and providing better protective function. Moreover, after the oral probiotic treatment, the abundance of *Lactobacillus* species/genus was found to be increased accordingly in some studies(De Alberti, Russo, Terruzzi, Nobile, & Ouwehand, 2015; Russo, Edu, & De Seta, 2018) but still lack of the identification at the strain level and the metastasis pathway is unknown. Whereas oral administration of lactobacilli could result in keeping a healthy vaginal microsystem or contribute to the vaginitis treatment (Ho et al., 2016). Two candidate strains selected in this study showed better survival ability under low pH and high concentration of bile salt conditions, indicating the possibility of oral probiotics for vaginal health.

Safety is an essential factor that should also be evaluated for the qualification of the putative probiotics. Bacteria have the potential to act as the host of antibiotic resistance genes, which can be transferred to pathogens after being taken. All two candidate strains were sensitive to almost all antibiotics except bacitracin and metronidazole, demonstrating the high-level safety of these strains and capable of being used adjunct with metronidazole, which is an antibiotic commonly used for the treatment of bacterial vaginosis. Also, fragmentized sequences for the tetracycline antibiotic gene in LG55-27 and TM13-16 were spotted by CARD annotation. What’s more, the genome-based annotation was also performed using VFDB, PHAST databases, no harmful genes hit harbored in strains LG55-27 and TM13-16, which illustrate that they are not human pathogens.

The proper formulation and dosing are the crucial factors for the effectiveness of probiotics. In this study, both oral and vaginal administrations were applied to assess the efficacy of isolates, so did different dosages and combinations. The results suggested a higher adhesion ability to the vaginal epithelium than intestinal epithelium, which may be due to the ignorance of assessing the strain adhesion ability to intestinal epithelium during the screening stage. In addition, the mixture of LG55-27 and TM13-16 displayed better improving effect than single strains. Moreover, we proved that both oral and vaginal use can improve the symptoms of vaginal infection. The high dose and low dose should be supplied vaginally and orally, respectively.

BV was associated with high levels of vaginal interleukin (IL)-1β and was downregulated after clindamycin treatment (Diaz-Cueto et al., 2009) The pro-inflammatory cytokines IL-1β and TNF-α were all significantly raised among women with STIs and/or BV. And after STI treatment, there were an overall reduction in that pro-inflammatory (Garrett et al., 2021). *L. johnsonii* inhibited the expression of pro-inflammatory cytokines such as IL-1β, TNF-α (Joo et al., 2011). The oral or vaginal administration of the *Lactobacillus* strain LG55-27 and TM-13 in BV rat model dampened the vaginal inflammation by decreasing the IL-1β and TNF-α, indicating the potential effectiveness of the strain for BV treatment. An interesting phenomenon occurred in this study, the levels of IL-1β with oral LG55-27 and TM13-16 administration increased. This case might be explained by the Jarisch Herxheimer reaction (Dhakal & Sbar, 2021), which was described an exacerbation of skin lesions in a syphilis patient after starting treatment with a mercurial compound. This likelihood was that the toxic metabolites produced by the pathogenic bacteria to resist the probiotic made response aggravates during the initial administration. The anti-inflammatory effect of oral administration might be due to its effects on immune responses through the gastrointestinal tract. Therefore, the response time was longer than vaginal administration and we couldn’t monitor the downward trend during the 6 days intervention. The multiple time points during the intervention and follow-up should be set up to monitor the dynamics of the inflammatory response.

## Acknowledgement

This research was supported by BGI Precision Nutrition (Shenzhen) Technology Co., Ltd. China, and the funding from Science and Technology Planning Project of Shenzhen Municipality (JCYJ20190809101409603). The authors declare no conflicts of interest.

## Figure and Table Legends

**Supplement Figure 1. The D-/L- lactate produced by two reference, candidate strain and its mix strain**.

**Supplement Table 1. The Score detail of the 98 *Lactobacillus* strains according to the BLAST result**.

