## Supplementary figures and images for "From whole genome to probiotic candidates: a study of potential *Lactobacillus* strain selection for vaginitis treatment"

### Supplement Figure 1

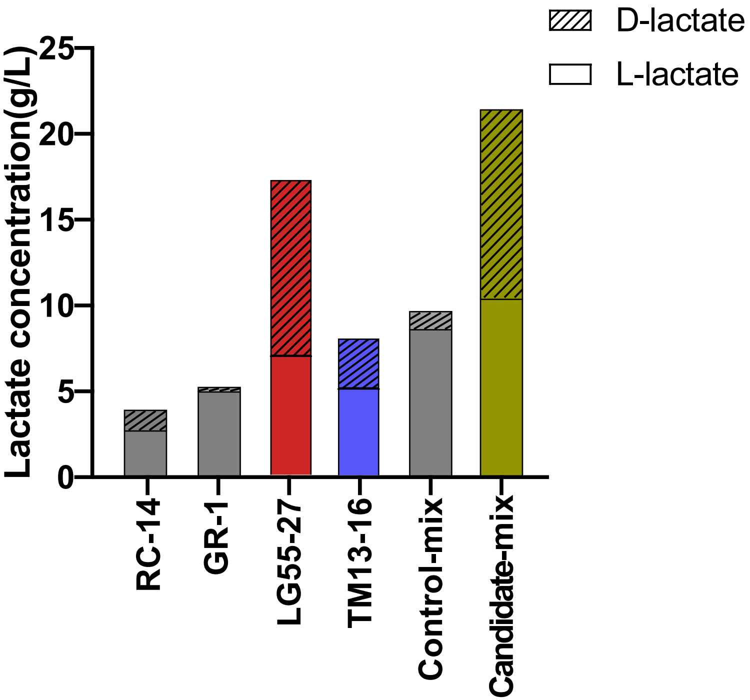
